# The integration of tandem gene repeats *via* a bacterial type-II toxin-antitoxin-mediated gene amplification (ToxAmp) system and stability visualisation in *Saccharomyces cerevisiae*

**DOI:** 10.1101/2024.01.30.578080

**Authors:** Samuel Evans, Zeyu Lu, Liam McDonnell, Will Anderson, Francisco Peralta, Tyson Watkins, Hafna Ahmed, Carlos Horacio Luna-Flores, Thomas Loan, Laura Navone, Matt Trau, Colin Scott, Robert Speight, Claudia E. Vickers, Bingyin Peng

## Abstract

Tandem gene repeats naturally occur as important genomic features and determine many traits in living organisms, like human diseases and microbial productivities of target bioproducts. Here, we develop a bacterial type-II toxin-antitoxin-mediated method to manipulate genomic integration of tandem gene repeats in *Saccharomyces cerevisiae* and further visualise the evolutionary trajectories of gene repeats. We designed a tri-vector system to introduce toxin-antitoxin-driven gene amplification (ToxAmp) modules, and accidentally re-visited the high-level capacity of multi-fragment co-transformation in *S. cerevisiae*. This system delivered the multi-copy gene integration in the form of tandem gene repeats spontaneously and independently from toxin-antitoxin-mediated selection. Inducing the toxin (RelE) expressing *via* a copper (II)-inducible *CUP1* promoter successfully drove the *in-situ* gene amplification of the antitoxin (RelB) module, resulting in ∼40 copies of a green fluorescence reporter (GFP) gene per copy of genome. The copy-number changes, increasing and decreasing, and stable maintenance were visualised using the GFP and blue chromoprotein AeBlue as reporters. Copy-number increasing happened spontaneously not depending on a selection pressure and was quickly enriched through toxin-antitoxin-mediated selection. In summary, the bacterial toxin-antitoxin systems provide a flexible mechanism to manipulate gene copy number in eukaryotic cells and can be exploited for synthetic biology and metabolic engineering applications.

**Table of Contents Graphic:** 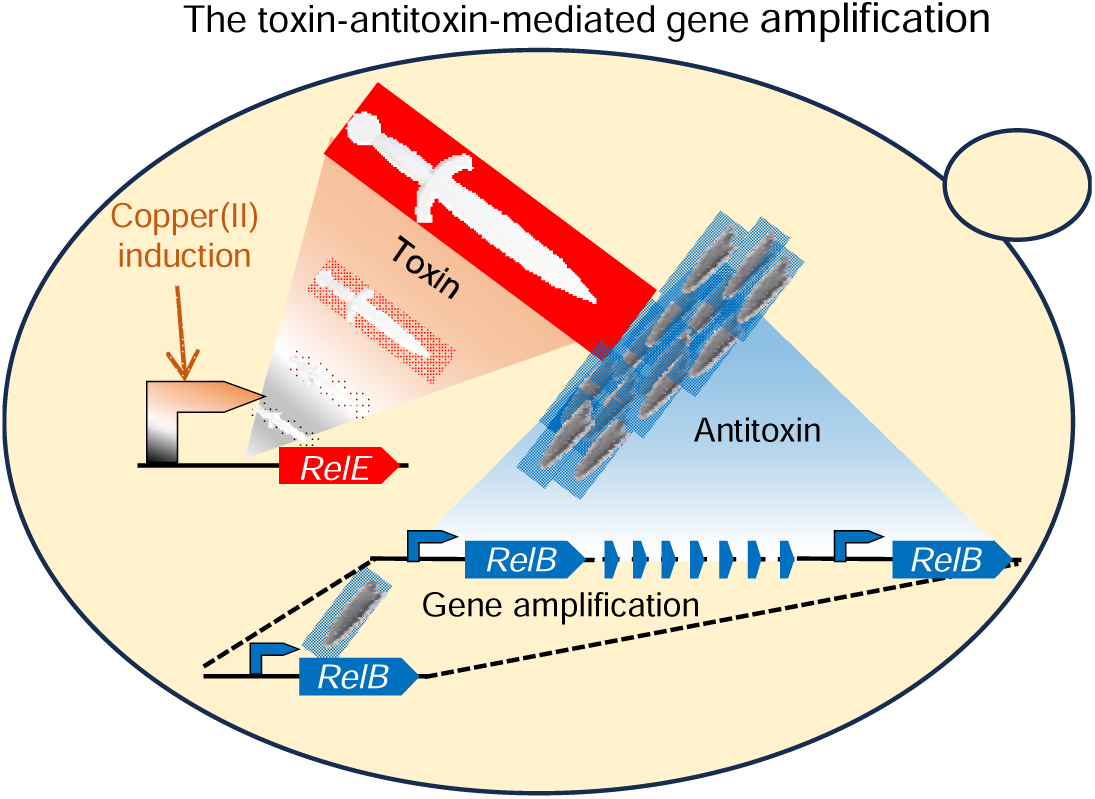

## Introduction

Tandem gene repeats, in the form of an array of two or more copies of a gene, commonly exist in cell genomes, i.e., ribosomal RNA genes (rDNA),^1^ oncogenic genes,^2^ drug resistant genes,^3, 4^ secondary metabolite biosynthetic genes,^5, 6^ and many other trait genes.^7^ Their occurrence is generally regarded as the result of gene amplification, which increases gene copy number (gene dosage) and provides a phenotypic advantage during natural or artificial selection.^3, 5^ Manipulating tandem gene repeats has been used for optimising bioproduction of small-molecular metabolites and proteins in metabolic engineering and synthetic biology.^8, 9, 10^ Examining tandem gene repeats is an important method in understanding the development of relevant genetic diseases in humans.^2^ Genetic manipulation methods have therefore been developed towards precise manipulation and examination of gene tandem repeats.^8, 11^

In the yeast *Saccharomyces cerevisiae*, tandem gene repeats have been investigated and engineered in a variety of studies. For example, rDNA repeats are used in understanding cell aging.^12^ Using the copper resistance gene *CUP1* gene as an example, the molecular processes for the formation of tandem gene repeats through gene amplification was interrogated.^13^ In metabolic engineering and synthetic biology, introduction of transgene tandem repeats is one of common engineering strategies to increase protein expression levels.^11, 14^ For example, we previously developed a haploinsufficiency-driven gene amplification method that delivered a tandem repeat array of up to ∼50 copies of transgenes in yeast chromosomes. This tandem-repeat gene integration is mitotically stable due to the selection pressure from a growth-determining haploinsufficient gene. Prior to this, it has been shown that the formation of a long tandem gene array in yeast genome through the yeast integrative plasmids.^15, 16^ In these plasmids, either an auxotrophic marker or a dominant resistant marker is used for plasmid maintenance or selection of clones with increased copy number.^10, 17, 18^ The tandem amplification of xylose and cellobiose utilization genes has also been seen during prolonged adaptive growth.^19, 20^ This adaptation mechanism is also linked to the growth benefits caused by increased gene dosages. Unintentional tandem-repeat integration of sesquiterpene synthetic genes was found in our rationally engineered yeast cell factories. In these sesquiterpene-producing yeasts, the variance of copy number likely occurred randomly rather than being driven by a strongly linked selection pressure.^21^ These show that gene amplification may happen as either a selection-force-driven process or a random process.

In addition to above conditions, gene stability and copy number can be controlled through a toxin-antitoxin system in bacteria and archaea.^22, 23^ The toxin-antitoxin systems are small genetic elements common in bacteria and archaea, comprising of two modules: a toxin module functioning as a killing or growth-inhibiting mechanism in the host cell and its antidote – antitoxin module.^23–25^ They were firstly discovered as a mechanism that stabilises mobile genetic elements,^26^ and were shown to possess many functionalities in maintaining genome stability,^18^ stress responses,^25, 27^ and pathogenicity.^28^ Eight types of toxin-antitoxin systems are currently described,^23, 24^ and the toxin-antitoxin pairs are known to be orthogonal.^25, 29^ Among them, the type II mRNA interferase toxin modules MazF and RelE have been introduced into yeast previously for counter selection, delivering inducible growth defects.^30, 31^ The toxic effects can be rescued through the expression of the antitoxin modules MazE and RelB.^31, 32^ The expression dosages of toxin RelE and antitoxin RelB were previously shown to control growth rate.^31, 33^ Therefore, exploiting toxin-antitoxin systems provides a potential a mechanism to tune the selection pressure for gene copy number manipulation in yeast.

In this study, we employed the type-II toxin-antitoxin RelE-RelB from *Escherichia coli* to drive the selection of tandem-repeat gene amplification. A tri-vector system was constructed to integrate the RelE module, the RelB module, and selectable marker into yeast genome. A copper (II)-inducible promoter was used to control the expression of RelE, which enables the copper (II)-inducible amplification of the RelB-linked construct. The RelB-linked construct carrying a yeast enhanced green fluorescent protein was amplified up to 40 copies per genome. The stability of the tandem gene repeats was further investigated. At the single-colony levels, the blue chromoprotein was used to visualise the trajectory of tandem gene repeats during yeast propagation.

## Results

### Multi-copy chromosome integration not depending on toxin-antitoxin selection

Upregulating RelE and downregulating RelB slow down yeast growth.^31, 33^ We hypothesized that adjusting expression balance between RelE and RelB might be used to generate a selection force to control gene amplification through formation of tandem gene repeats. For example, if RelE were to be expressed at a relatively high level and RelB was expressed at a relatively low level, there would be a selection pressure for cells in which the RelB expression cassette had been duplicated; i.e., the increased titre of RelB caused by gene duplication would alleviate the growth defect caused by high titres of RelE.

To test this hypothesis, we designed a three-plasmid genetic system to introduce the expression cassettes of RelE, RelB, an hygromycin B resistant marker (Hph), and a yEGFP reporter (**Figure 1**). The expression cassettes of RelE and Hph are closely linked and can be integrated into the yeast chromosome through the homologous recombination targeting to the *HO* locus. The expression cassettes of RelB and yEGFP are closed linked and can be integrated into *HO* promoter region to form two repeated *HO* promoters. In the first design, RelE is under the control of a strong constitutive promoter (the *TDH3* promoter), and RelB is under the control of a medium strong promoter (the *RPL8B* promoter; **Supplementary Figure 1**). The *TDH3* promoter is ∼six-fold stronger than the *RPL8B* promoter ^34^. The transformation efficiency for this design was quite low. In one of many attempts, two large colonies grew with only one showing a strong green fluorescence, and in other attempts, transformation was not successful. The low transformation efficiency might be as the result of acute toxicity caused by RelE overexpression and the RelB module was not efficiently amplified to rescue the toxicity during the transformation selection process.

**Figure 1.**
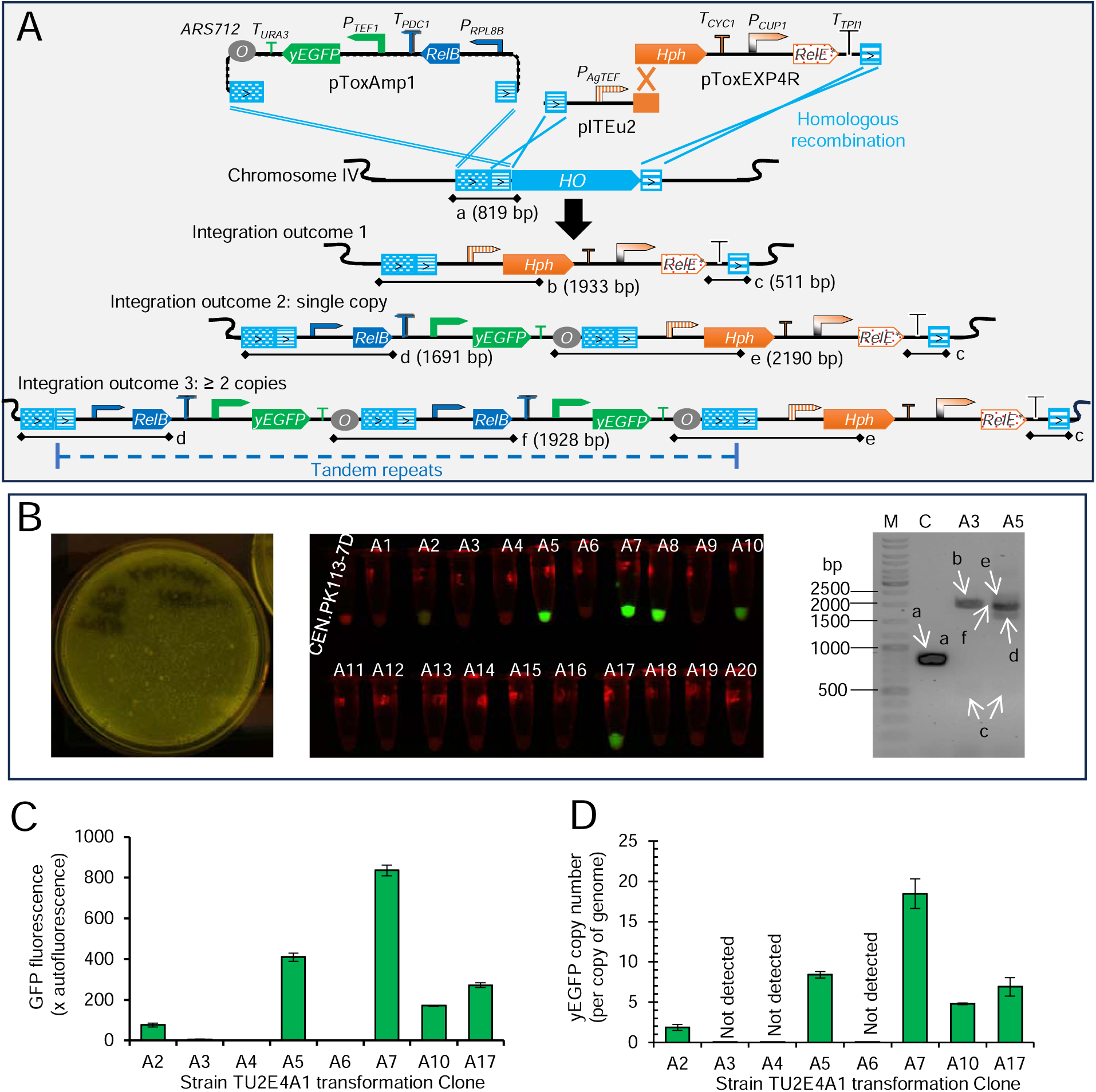
A tri-vector system to deliver the integration of tandem gene repeats, not depending on the functional toxin-antitoxin selection. **A**. Diagram of the tri-vector systems and integration outcomes. Fragments from the three vectors: pToxAmp1, pITEu2, and pToxEXP4R. ARS712, replication origin. *T_XXXN_*, the terminator of *XXXN* gene. *P_XXXN_*, the promoter of *XXXN* gene. yEGFP, yeast enhanced green fluorescent protein. RelB, antitoxin gene, RelE*, mutated non-functional (‘dead’) toxin gene. *HO*, YDL227C gene from *S. cerevisiae*. a-f, PCR fragments for genotype determination. **B**. Transformation outcomes for strain TU2E4A1: a transformation plate under the blue-light illumination, the yeast cells of isolated clones under blue-light illumination, and colony PCR results. M, DNA ladder marker (Thermo Fisher #SM0331); C, the control strain CEN.PK113-7D. A1-A20, isolated clone number. a-f, fragments in **A**. **C** & **D**, yEGFP fluorescence and gene copy numbers in selected clones. Mean value ± standard deviation are shown (n = 3 technical replicates). yEGFP fluorescence was determined using Accuri C6 flowcytometry.

In the second design, we used a copper(II)-inducible promoter, the *CUP1* promoter, to control the RelE expression (**Figure 1**). The *CUP1* promoter exhibits a weak basal activity in the absence of additional copper(II),^34^ which may allow the positive selection of the integration of the RelB-yEGFP module. In the presence of additional copper(II) (e.g., 100 μM), the *CUP1* promoter activities can be induced to the levels ∼three-fold stronger than the *RPL8B* promoter activities.^34^ In theory, this could provide a mechanism to induce RelE expression for selection of the amplification of the RelB-yEGFP module. We transformed strain CEN.PK113-7D with the constructs of this design,^35^ and transformants were selected on the yeast extract-peptone-dextrose (YPD) hygromycin B plates without copper(II) addition. The fluorescence in most of colonies was not detectable (**Figure 1**).

Twenty colonies were picked and grown them overnight in liquid culture. The resulting yeast cells were collected and washed. Five out of twenty colonies showed green fluorescence (**Figure 1**). We characterised the yEGFP fluorescence using flow cytometry and copy number using quantitative real-time PCR in these five colonies and three control colonies that did not show increased fluorescence (**Figure 1**). GFP fluorescence and gene were not present in the control colonies (**Figure 1**). In the five colonies, the copy number of yEGFP gene ranges from one to ten, and GFP fluorescence was positively correlated with yEGFP gene copy number (*R*^2^ = 0.99; **Figure 1**; **Supplementary Figure 2**).

We performed yeast colony PCR to verify the integration of transgenes in the three-plasmid system. In the fluorescent clone A7, the amplification of bands d, and f, indicates the integration of yEGFP-RelB construct at *HO* promoter locus, the band e for the formation of tandem repeats, and the bands c and f for the integration of the RelE construct (**Figure 1A&B**). Multi-copy integration in clone A7 was confirmed through Oxford nanopore whole genome sequencing (**Supplementary Figure 3**). In the clone without the yEGFP fluorescence, the amplification of band b indicates the integration of the *RelE* construct (**Figure 1A&B**). This was unexpected. These fluorescent colonies were grown in the YPD medium with addition of 100 μM copper (II) sulfate from a very low OD_600_ (∼0.004) and transferred the cultures under the same conditions. The fluorescence levels did not increase. Later, we found that there was a single base pair insertion after the second codon of *RelE* open reading frame (**Supplementary Figure 4**). This indicates that the toxin *RelE* was not functioning in selection of the multi-copy integration of the yEGFP-RelB construct. Therefore, we presume that the integration of the yEGFP-RelB construct happened as a co-transformation event that coupled with the transformation of the selectable marker Hph and “dead” (inactive)-RelE constructs. Together, these data indicate that multi-copy chromosome integration in these strains does not depend on the toxin-antitoxin selection.

### Toxin-antitoxin-driven gene amplification

Although the multi-copy chromosome integration does not depend on the toxin-antitoxin selection, it is still interesting to see whether increasing the toxin expression can provide an evolutionary pressure to drive or select the amplification of antitoxin-expressing module. We therefore attempted to reconstruct the plasmid with the RelE gene under the control of the *CUP1* promoter but failed to generate a plasmid with the correct sequence. We presume that the *CUP1* promoter might be active in *E. coli*. To solve this problem, we inserted the *RPL28* intron in the middle of the RelE gene. We successfully constructed two RelE_[*RPL28* intron]_ constructs: in one, the *CUP1* promoter was used to control RelE expression, and in the other, the *GAL1* promoter was used (**Figure 2A**).

**Figure 2.**
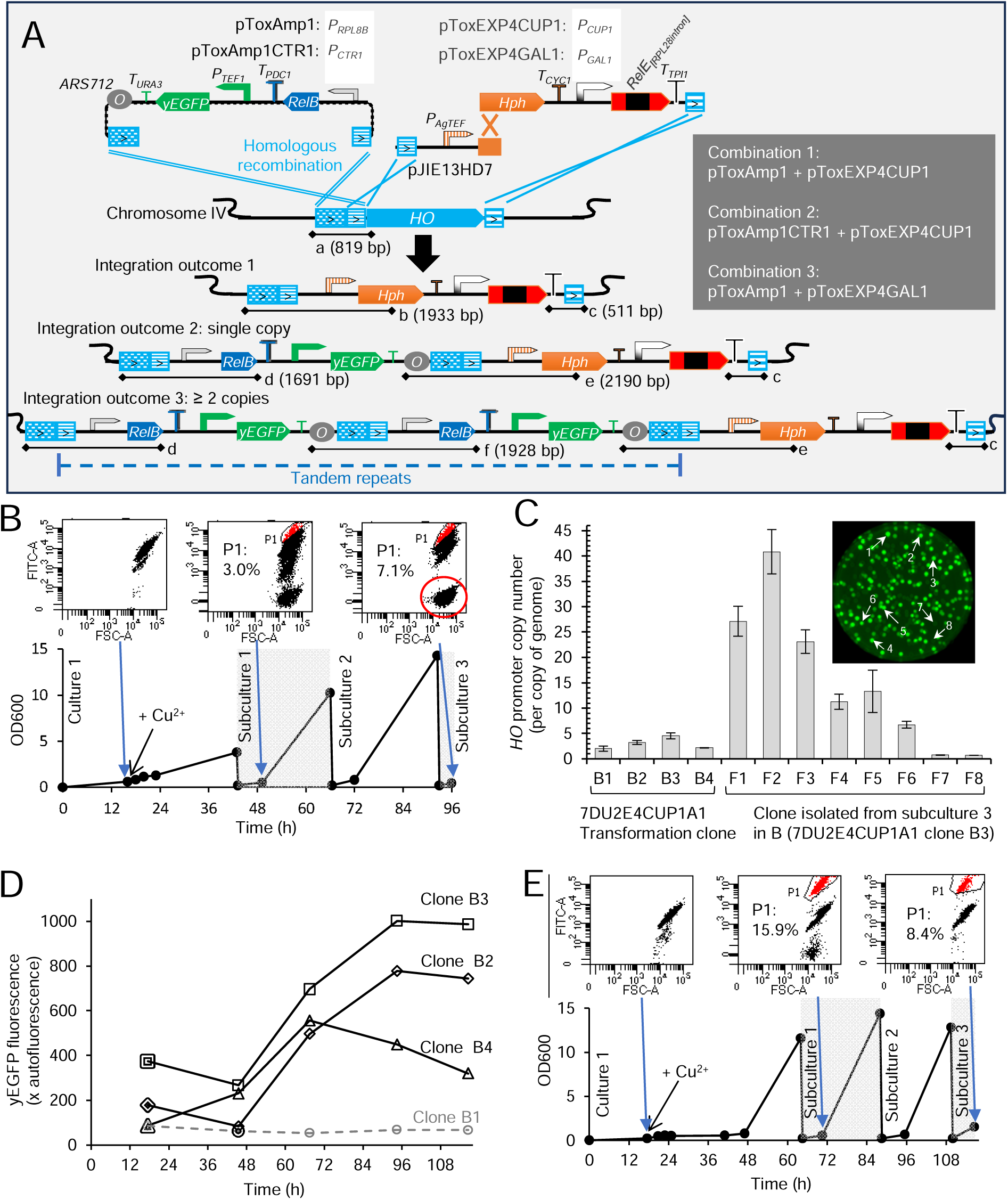
The copper (II)-inducible *in vivo* gene amplification driven by the bacterial type-II toxin-antitoxin (RelE-RelB) pair (ToxAmp). **A**, Diagram of the tri-vector systems for ToxAmp and integration outcomes. Fragments from the three vectors: pToxAmp1/pToxAmp1CTR1, pJIE13HD7, and pToxEXP4CUP1/pToxEXP4GAL1. ARS712, replication origin. *T_XXXN_*, the terminator of *XXXN* gene. *P_XXXN_*, the promoter of *XXXN* gene. yEGFP, yeast enhanced green fluorescent protein. RelB, antitoxin gene, RelE_[RPL28intron]_, the toxin gene RelE inserted with the *RPL28* intron. *HO*, YDL227C gene from *S. cerevisiae*. a-f, PCR fragments for genotype determination. **B**, The evolutionary effects of copper(II) induction in Combination 1 transformant (7DU2E4CUP1A1) clone B3 (Supplementary Figure 5; the isolate was not pure). **C**, *HO* promoter copy numbers in Combination 1 transformants (7DU2E4CUP1A1: B1-B4) and the isolates (F1-F8) from the evolved culture. **D**, Copper(II)-induced evolutionary effects on overall yEGFP fluorescence for the pure Combination 1 transformant isolates (B1-B4). The markers with double lines indicate the starting time of copper(II) induction. **E**, Copper(II)-induced evolution for the pure Combination 1 transformation isolate B4, harbouring a single-copy integration of pToxAmp1. Synthetic minimal media, yeast nitrogen base wo/amino acids + glucose, were used for yeast cultivation. Copper(II) sulphate of 100 nM was added to induce the evolution. Mean value ± standard deviation were shown (C; n = 3 technical replicates). B & E, FITC PMT voltage = 350. D, n = 1. The details of evolutionary cultivation for pure clones B1-B3 are shown in **Supplementary Figure 9**.

In the antitoxin RelB-yEGFP module, the *RPL8B* promoter was used to control RelB expression. We also made another RelB-yEGFP constructs, in which the copper-repressible *CTR1* promoter was used to control RelB expression (**Figure 2A**).

We transformed CEN.PK117-7D with RelE_[RPL28 intron]_-expressing module and RelB-yEGFP-expressing module in three combinations: *P_CUP1_*-RelE with *P_RPL8B_*-RelB (combination 1), *P_CUP1_*-RelE with *P_CTR1_*-RelB (combination 2), and *P_GAL1_*-RelE with *P_RPL8B_*-RelB (combination 3). Transformants were selected on YPD agar supplemented with hygromycin. Both fluorescent clones and non-fluorescent clones were obtained. Fluorescent clones included bright clones and dim clones (**Supplementary Figure 5**). Yeast colony PCR results indicate that the dim clones have a single-copy integration of the RelB-yEGFP module, the bright clones have a multi-copy integration, and the non-fluorescent clones have no integration of RelB-yEGFP-expressing module (**Supplementary Figure 5**).

For each transformation combination, we chose two clones with a single-copy integration and two clones with a multi-copy integration to test whether the induction of toxin RelE expression could drive and select for amplification of the RelB-yEGFP module. For the clones with the *P_CUP1_*-RelE and *P_RPL8B_*-RelB cassettes (Combination 1), cells were grown in MES-buffered YNB-glucose broth with 100 nM copper (II) sulphate. For the clones with the *P_CUP1_*-RelE and *P_CTR1_*-RelB cassettes (Combination 2), cells were grown in YPD broth with 100 nM copper(II) sulphate, as they did not grow in YNB-glucose broth. For the clones with *P_GAL1_*-RelE and *P_RPL8B_*-RelB cassettes (Combination 3), the cells were grown in MES-buffered YNB-galactose broth. Only cells with the *P_CUP1_*-RelE and *P_RPL8B_*-RelB cassettes (Combination 1) had populations with increased yEGFP fluorescence were observed (**Figure 2B**). In the cells of other Combinations, we did not observe the evolution towards increased yEGFP fluorescence.

We investigated the copy numbers of the Combination 1 (*P_CUP1_*-RelE and *P_RPL8B_*-RelB) clones before the copper(II)-induced evolution. Consistent with the colony PCR results (**Supplementary Figure 5**), clone B1 and clone B4 harboured single-copy integration (two copies of *HO* promoters), and clone B2 and clone B3 harboured two or four copies of integration.

We isolated the single colonies from the evolved culture of clone B3. Colonies showing different levels of GFP fluorescence were isolated. The copy number increased up to ∼40 copies per copy of genome (**Figure 2D**). In the clones with very weak green fluorescence, the *HO* promoter copy number was ∼one (**Figure 2D**). Integration of a single copy of the GFP-RelB construct resulted in two copies of the *HO* promoter per copy of genome. This indicates a lack of the integration of *GFP*-RelB construct in these clones.

To understand what happened in the cells that lost the fluorescence during evolution, we performed the whole genome sequencing for Combination 1 strain clone B4 (the single-copy integration clone; **Supplementary Figure 5**) and its evolution culture. Genome assembly confirms the single-copy integration in Combination 1 strain clone B4 (**Supplementary Figure 6**) and shows the wildtype *HO* (*YDL227C*) locus, indicating a contamination in the evolution culture (**Supplementary Figure 7**). The contamination was likely caused by a procedural mistake: the colonies picked from the transformation plates were replicated on the YPD agar in the absence of hygromycin selection (**Supplementary Figure 5**). The sequencing coverage was low for the clone B4 evolution culture sample, which prevented the assembly of the RelE module and the RelB-yEGFP module; but we could find the nanopore reads to confirm their existence in this sample (**Supplementary Figure 8**).

To test the evolutionary outcome of the toxin-antitoxin-driven gene amplification, we isolated the single colonies for the clones of Combination 1 and Combination 3 strains and re-performed the copper(II)-induced evolution (**Figure 2D & 2E, Supplementary Figure 9 & 10**). Induction of RelE expression resulted in a population of cells without yEGFP fluorescence, but this population of cells disappeared in the next round of subcultures (**Figure 2E, Supplementary Figure 9A**-C **& 10B-E:** in seven out of eight experiments). The clones without RelB-yEGFP modules may escape from RelE toxicity after prolonged cultivation (**Supplementary Figure 9D-E & 10F-G**). Three out of four clones for the Combination 1 strain, two multi-copy integration clones and one single-copy integration clone, showed a copper(II)-induced increase in the yEGFP fluorescence (**Figure 2D**). One out of four clones for Combination 3 strain, the single-copy integration clone, showed a small population having yEGFP fluorescence increased upon galactose induction of RelE expression (**Supplementary Figure 10C: P1, 0.6%**).

### Fluorescent reporter for stability visualisation of tandem repeat sequences with toxin-antitoxin selection

Copper(II)-mediated induction of RelE expression triggered the toxin-antitoxin-driven gene amplification, which delivered the multi-copy integration of heterologous genes on the chromosome (**Figure 2C**). This process often occurs during the process of DNA replication and repair,^36^ but the resulting tandem-repeat structures may not be mitotically stable. We were interested in the stability of the resulting multi-copy integration. Many techniques and statistical methods have been used to characterise the stability of tandem repeats in yeast.^37^ Characterisation methods including the examination of gene copy number using quantitative real-time PCR (such as in Figure 1D and Figure 2C), the evaluation of selectable marker stability through growth testing, and whole-genome sequencing.^8, 20^ However, these methods can only evaluate a limited number of samples. Alternative methods may be used for visualising the evolution trajectory of tandemly repeated gene locus. In our test, we observed that yEGFP fluorescence was well correlated with its gene copy number (Supplementary Figure 2). Therefore, yEGFP fluorescence can be used as a proxy for copy number to examine the stability.

To test the stability, we sequentially sub-cultured the six multi-copy integration clones, isolated after copper(II)-induced gene amplification (Figure 2C: F1-F6). The six clones were continuously sub-cultured in four media: YPD broth, YPD broth supplemented with 100 nM copper(II), YNB-glucose broth, and YNB-glucose broth supplemented with 100 nM copper(II). The stability profiles in YPD broth were not influenced by the presence of copper(II) (**Supplementary Figure 11A, 11B & Figure 3A**). Clones F1, F2, and F3, harbouring > 20 copies of the integrated region, decreased in the average GFP fluorescence over the subcultivations (**Figure 3A**). At the population level, the proportion of the cells having high-level GFP fluorescence decreased, and the proportion of the cells having low-level GFP fluorescence increased (**Figure 3B**). Clones F4, F5, and F6, harbouring 6-15 copies of integration, were relatively stable, but also showed a slow trend towards losing the population of highest-level fluorescent cells (**Figure 3A and 3B**).

**Figure 3.**
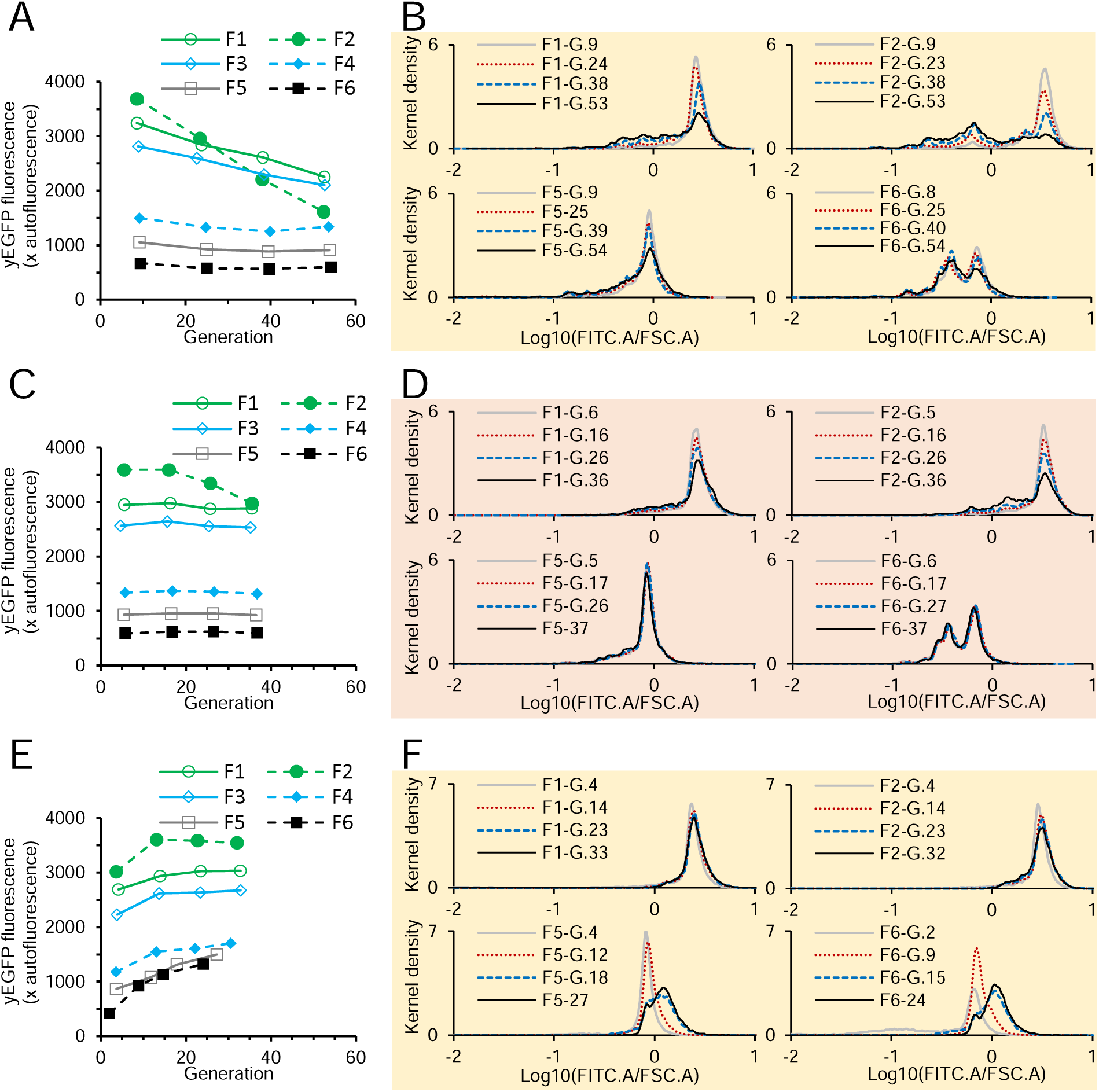
The stability of the multi-copy integration of the yEGFP expression cassettes *via* the *RelE-RelB*-driven gene amplification. (**A** & **B**) YPD broth supplemented with 100 nM copper(II). (**C** & **D**) YNB-glucose broth. (**E** & **F**) YNB-glucose broth supplemented with 100 nM copper(II). G., generation. A-F, n = 1. The bandwidth of kernel density is 0.01.

In YNB-glucose broth, the multi-copy integrations showed a better stability than those in YPD or YPD + Copper(II) broth (**Figure 3A-D**). However, for the clones harbouring > 20 copies of the integration, the proportion of high-level fluorescent cells decreased at a rate similar to those growing in YPD-based broth (Figure 3B and 3D: F1, F2; Supplementary Figure XC-D: F3). For the clones harbouring 6-15 copies of the integration, the cell population did not change dramatically.

In YNB-glucose + Copper(II) broth, the population of the cells having > 20 copies of the integration was relatively stable (Figure 3E and 3F; Supplementary XE). The clones having 6-15 copies of the integration evolved towards increased yEGFP fluorescence, consistently with the toxin-antitoxin-driven gene amplification.

### Chromoprotein for stability visualisation of tandem repeat sequences

Using yEGFP as the reporter, we examined the stability of tandem repeat sequences in liquid culture (Figure 3). At the single-colony levels, we also observed the potential instability of yEGFP-expressing cells, shown as the colonies patterned with sectors under blue-light illumination (Supplementary Figure 1). We were interested in developing an educational kit to demonstrate the phenomenon of genetic stability of tandem repeat sequences. For this purpose, a chromoprotein reporter might be better than a fluorescent reporter, since it can be observed with the naked eye. Therefore, we designed and synthesised a pToxAmp12 construct (Figure 4A), in which the yEGFP expression cassette was replaced with an expression cassette of the blue chromoprotein from *Actinia equina* (AeBlue).^38^ In the AeBlue expression cassette, a strong constitutive *TDH3* promoter and a synthetic mini-terminator (T*_synth_*_7_)^39^ were used to control AeBlue expression. However, this experiment was performed before the best construct for the toxin-antitoxin-driven gene amplification (Figure 2A) was revealed, hence we used a non-optimal design. We used the constructs with the *RelB* under the control of the *CTR1* promoter (pToxAmp12: Figure 4A) and the construct with the dead-RelE (pToxEXP4R: Figure 1A & Figure 4A). The three constructs were transformed into *S. cerevisiae* strain CEN.PK113-7D. Blue colonies appeared together with the white colonies on the selection plate, indicating the co-transformation of *AeBlue-RelB* construct with the *Hph* and dead-*RelE* constructs.

**Figure 4.**
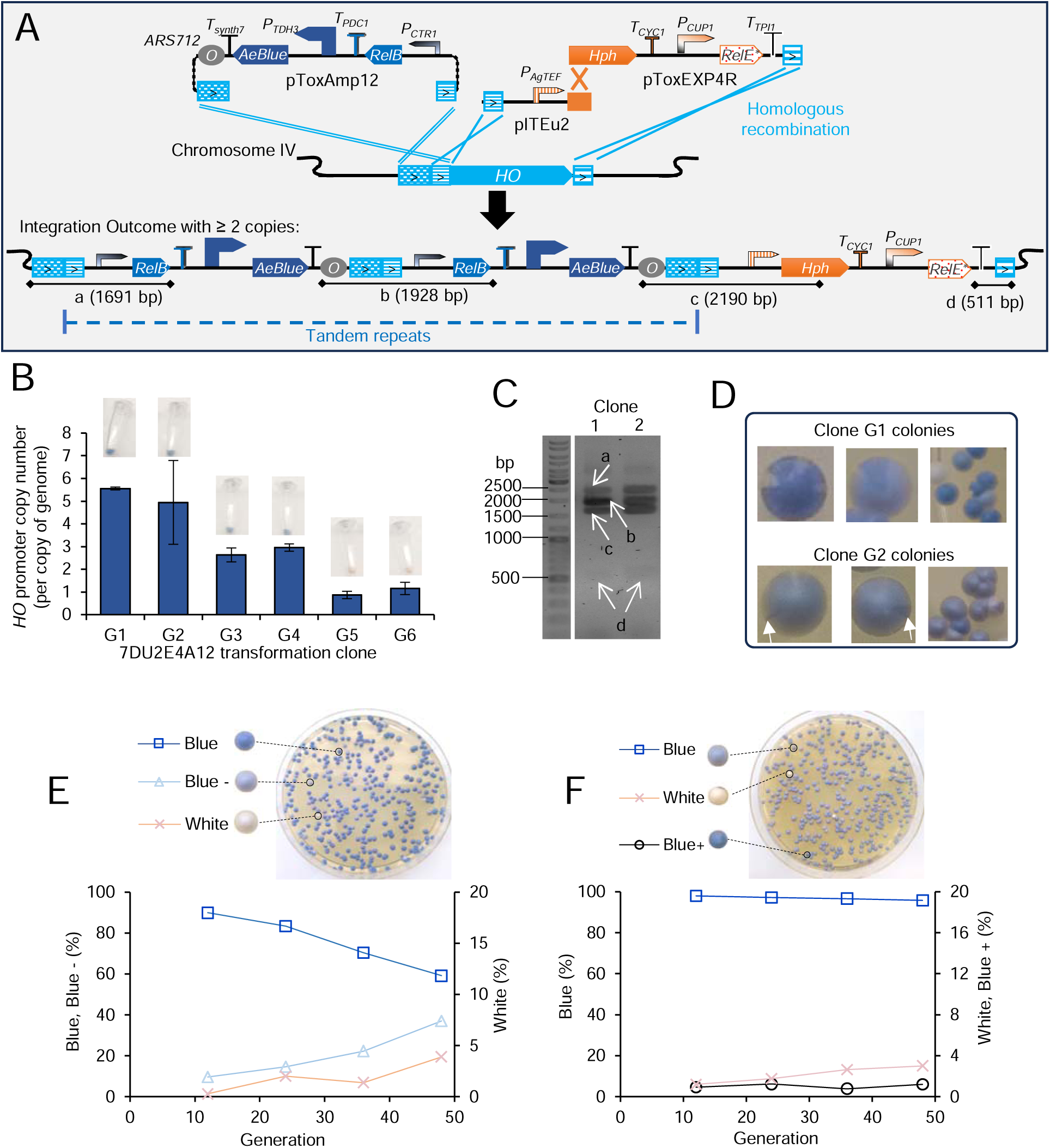
Using AeBlue to visualise the stability of tandem gene repeats in yeast. (**A**) Genetic constructs for introduction of multi-copy integration of AeBlue overexpression cassette. (**B**) *HO* promoter copy number in the yeast clones isolated from transformation plates. (**C**) Yeast colony PCR to verify the integration pattern. (**D**) Single yeast colonies showing the stability of multi-copy integration of AeBlue overexpression cassette. White arrows show the sectors with increased saturation, which means increased copy number. (**E** & **F**) Stability analysis *via* counting the colonies with three-scaled hue in continuous transferring cultivation: G1 (**E**) and G2 (**F**). Blue, the colonies showing the initial blue hue. Blue -, the colonies showing decreased blue hue. Blue + the colonies showing increased blue hue. White, the colonies appearing in pale white colour. Mean values ± standard deviation are shown in B (n = 3 technical replicates). (E & F), n = 1.

We isolated four blue clones (**Figure 4B. Clones G1, G2, G3, and G4**) and two white clones (**Figure 4B. Clones G5 and G6**). The colony PCR results support the multi-copy integration of the AeBlue-*RelB* construct in the blue clones. We used quantitative PCR to analyse the copy number of the *HO* promoter region that was amplified together with AeBlue and RelB. This further confirmed that the multi-copy integration was not dependent on toxin-antitoxin selection. In the white clones, the *HO* copy number is one, indicating that the AeBlue-RelB construct was not integrated. In the blue clones, the *HO* promoter copy-number ranges from three to six, indicating multi-copy integration of the AeBlue-RelB construct. Oxford Nanopore whole genome sequencing confirmed the three-copy integration of the AeBlue-RelB construct at the *ho* locus in Clone G2. Clone G1 (five-copy integration) shows a darker blue colour than clones G2/G3/G4 (two-copy integration). This suggests that the colour saturation of the yeast colonies correlate with the copy numbers of the AeBlue gene. This allows a direct observation of the stability and variation of AeBlue tandem repeats in prolonged cultivation.

We sub-cultured two blue clones (**clone G1 and clone G2**) over ∼48 generations. Yeast cells were spread on YPD plates for each cultivation and incubated at room temperature for a week for full development of AuBlue hue. The colonies on the plate did not show a homogenous blue colour. Clone G1 colonies showed patterned sectors with decreased hue for most cases (**Figure 4D**). In Clone G2, sectors with increased hue appeared in very few cases (**Figure 4D**).

To evaluate the strain stability, we counted the colonies into three scales of blue hue (**Figure 4E & 4F**). For Clone G1, the proportion of the original blue colonies decreased to ∼59% over 48 generations, and less-blue colonies (Blue -) and white colonies gradually increased (**Figure 4E**). For Clone G2, having lower copy number of AeBlue construct and less blue hue than Clone G1, ∼96% colonies maintained the parental blue colour and ∼ 1 % colonies showed increased blue colour (Blue +). In the clone G2 subculture, a brown colony appeared on the agar plate. Subsequent characterisation showed that a deficiency of adenine synthesis led to the production of orange dye and the multi-copy integration can be lost in 10% cells after 45-generation sub-cultivation (**Supplementary Result 1**). The mixture of orange pigment and blue pigment may generate the brown colour.

## Discussion

To manipulate the tandem gene repeats in *S. cerevisiae*, we developed a bacterial type-II toxin-antitoxin-mediated gene amplification (ToxAmp) method in this work. The integration of the tandem-repeat construct does not depend on the toxin-antitoxin-mediated selection. In the ToxAmp method, inducing the expression of toxin RelE can be detrimental, but it can be rescured by an increase in gene dosage of the antitoxin RelB module. The tandem gene repeats are not stable; increase and decrease of copy number happen spontaneously, and the toxin-antitoxin-mediated selection pressure is important to maintain the high-copy stability.

Co-transformation of two different plasmids into *S. cerevisiae* were firstly reported in 1970s.^40^ We have co-transformed four plasmids to complement four auxotrophic selection markers to establish metabolic engineering strains in one step.^41^ Co-transformation of an episomal plasmid and a genome-integrating DNA fragment was also reported in the 1980s, but the co-transformation ratios were low (< 2 %).^42^ Here, we accidentally re-visited the co-transformation event in *S. cerevisiae* through the genome-integrating tri-vector system (Figure 1). The co-transformation efficiency was higher, ∼ 30 %, which might be due to the specific design for the homologous fragments (**Figure 1**). This is consistent with *S. cerevisiae*’s high-capacity of homologous recombination-based *in vivo* DNA assembly, which was first reported in 1980s and only well exploited for large-DNA fragment assembly 20 years later. ^43^

Multi-copy integration methods were previously investigated in *S. cerevisiae*, and other yeasts for applications in metabolic engineering.^8, 44^ They include high-concentration antibiotic-based selection ^14, 16, 44, 45^ and ribosomal DNA or delta-sequence-targeting integration.^46^ Recently, we showed that the typical yeast integrative plasmid (YIp) could be used to deliver multi-copy genome integration in the pattern of tandem gene repeats.^21^ We note that YIp delivering multi-copy genome integration has been previously reported in 1981.^47^ The current work further confirms that the multi-copy genome integration feature happens during the transformation, and does not depends on a specific selection pressure (**Figure 1 & Figure 4**). These findings may help to leverage engineering capacity to use the existing yeast integrative vectors, which were not originally designed for multi-copy genome integration, to select the multi-copy integration and explore the phenotypic advantages.

Using AeBlue as the reporter, we observed again that gene amplification happened spontaneously, and does not depending on the selection pressure (**Figure 4D & 4F**). The induction of RelE under the control of the *CUP1* promoter provided the selection to enrich the cells with increased RelB copies (Figure 3, Supplementary Figure 6). But the acute RelE toxicities *via* the expression under the control of the *GAL1* promoter, two-fold stronger than the *CUP1* promoter under inductive conditions,^34^ caused growth arrest (**Supplementary Figure 10**). In these cases, other unknown mechanisms were adopted to escape RelE toxicities (**Supplementary Figure 10**). A recent report showed that that spontaneous mutagenesis inactivated RelE selection in *Pseudomonas protegens* and translational pairing between *RelE* and an isoleucine synthetic enzyme, *ilvA,* overcame the selection of inactivation mutations.^48^ Consistent with this, reduced ilvA/RelE was found to contribute to a better selection effect. Here, we observed that the basal expression of *RelE* under the control of the *CUP1* promoter was sufficient to cause growth arrest in YNB broth without 100 nM copper(II) supplement (**Supplementary Figure 6D and 6F**). These data suggest that strong expression is not essential to drive ToxAmp.

The tandem gene repeats did not show a complete mitotic stability, shown as the decreasing in over-all yEGFP fluorescence and the cells expressing the high-level yEGFP (**Figure 3**) and the appearance of the colonies and the colony section showing little or no expression of AeBlue protein (**Figure 4**). In both yEGFP and AeBlue cases, the lower-copy integration was more stable than the higher-copy integration (**Figure 3 & Figure 4**). Meanwhile, selection pressure is important to maintain the stability of high-copy integration. In YPD broth, 100 μM copper(II) was not sufficient to induce the toxin-antitoxin-mediated selection, which might be due to the copper(II)-chelating properties of amino acids.^49^ It has been shown that the copper(II) concentration needs to be in the mM range to induce the *CUP1* promoter.^50, 51^ In YNB broth, the integration of 6-15 copies can be stably maintained. This might be due to the basal expression of the *CUP1* promoter in YNB media, ^34^ whereas the *CUP1* promoter is not active in YPD broth without copper(II) addition.^51^ In the ToxAmp construct (Figure 2), the *RPL8B* promoter is used to control the expression of antitoxin *RelB*, and the *CUP1* promoter for toxin *RelE*. The strength of the *RPL8B* promoter controlling *RelB* expression is 6.5-fold higher than the basal strength of the *CUP1* promoter.^34^ This suggests that the antitoxin should be expressed at the level much stronger than the toxin to reach the proper balance for detoxification.

In summary, we observed the randomness of copy-number change in tandem gene repeats, including copy-number increases and copy-number decreases. The change does not depend on the selection pressure. However, transcriptionally active genes (*P_TEF1_-yEGFP* and *P_RPL8B_-RelB*) and a late-firing autonomous replicative sequence ARS712 may stimulate the gene amplification in the ToxAmp design.^13, 52^ In the absence of the selection pressure, the cell population harbouring high-copy-number repeats rapidly collapsed. In contrast, selection pressure could rapidly enrich for the high-copy-number integration at population levels. These observations indicate the challenges in cancer treatment through inhibition on the gene product of oncogenic tandem repeats ^53^ and the prevention of the relevant human diseases,^54^ as tandem repeats inevitably occur during natural proliferation processes. The AeBlue-expressing ToxAmp constructs provide an eukaryotic system to visualise the trajectories of tandem gene repeats in complement to the *E. coli* system,^55^ and can be used for educational purposes to demonstrate genetic variations in cell proliferation. Introduction of stable multi-copy gene integration can be used for optimisation of bioproduction in metabolic engineering,^8^ and many bacterial type-II toxin-antitoxin pairs can be further exploited.^23, 24, 56^

## Materials and methods

### Plasmid and yeast strains

Plasmids and *S. cerevisiae* strains are listed in Supplementary Data file 1. Plasmid sequences are listed in Supplementary Data file 2. Molecular Cloning Designer Simulator was used to simulate plasmid construction.^57^

### Flow cytometry

Yeast cells were grown in YNB media containing 6.9g L^-1^ yeast nitrogen base without amino acids (YNB, FORMEDIUM#CYN0402) and 20 g L^-1^ glucose to the exponential growth phase (OD_600_IJ≤IJ2) for characterisation of the yEGFP fluorescence. A BD Accuri™ C6 flow cytometer (BD Biosciences, USA) was used for fluorescence analysis in single cells (**Figure 1**). A forward-scatter height (FSC.H) threshold of 250,000 was used to exclude cellular debris particles. A 488IJnm laser was used to excite GFP fluorescence. The detector equipped with a 530/20 bandpass filter was used to monitor the fluorescence (FL1.A) and 10,000 events were recorded for each sample. BD Csampler software (BD Accuri C6 software version 1.0.264.21) was used to determine mean values of forward-scatter area (FSC.A), side-scatter area (SSC.A) and FL1. A.

A BD FACSCanto II flow cytometer (Firmware version 1.49) was used to measure the yEGFP fluorescence (**Figure 2** and **Figure 3**). A 488 nm laser was used to excite the yEGFP fluorescence, and yEGFP fluorescence was monitored at FITC-A channel through a 530/30 nm bandpass filter between two long-pass dichroic mirrors of 556 nm and 502 nm. FITC-A channel PMT voltage was adjusted accordingly to make the FITC-A channel measurement in the range of detection (the value = 300, 350, or 552). A BD FACSDiva™ Software (version 8.0.1; CST Version 3.0.1; PLA version 2.0) was used extract the mean values of FITC-A, forward scatter (FSC-A, PMT voltage = 200; gating threshold = 3000), and side scatter (SSC-A, PMT voltage = 200).

The fluorescence level of GFP was expressed as the fold of the background fluorescence in CEN.PK 113-7D cells grown to the exponential growth phase according to the previously published equation.^34^ To evaluate the kernel density, the data in .fsc file was saved into .csv file using an online converter (https://floreada.io/fcsconvert), and the data were extracted and analysed using R (Supplementary code 1).

### Quantitative real time PCR

NEB Luna^®^ Universal qPCR Master Mix was used, and PCR reactions were prepared in accordance with the manual. Genomic DNA for the reference and experimental samples were diluted with nuclease free water to 0.1 ng µl^-1^ and used as templates in quantitative real-time PCR. Primers for quantitative real-time PCR for genes *ACT1*, *yEGFP*, and the *HO* promoter (Supplementary Data 1). Real-time PCR reactions were run in a 384-well plate using a Bio-Rad CFX384™ Touch Real-Time PCR Detection System. SYBR-green fluorophore was used in analysis. Genomic DNA extracted from strain ILHA Y2A-GFP ^58^ was used for preparing the standard curves and used as the single-copy reference for the yEGFP gene and the *HO* promoter. The gene copy numbers were calculated using the 2^-ΔΔCt^ method.^59^

### Prolonged transferring cultivation

#### Induced ToxAmp cultivation

For Combination 1 clones (Figure 2 and Supplementary Figure 6), cells were inoculated to OD600 of 0.008 in YNB-glucose broth. Copper(II) sulphate was added to a final concentration of 100 µM when OD600 was above 0.2. In the following subcultures, YNB-glucose broth supplemented with 100 µM copper(II) sulphate was used and cells were inoculated to OD600 of 0.2 to start a new culture. For Combination 2 clones (Figure 2), YPD broth was used instead of YNB-glucose broth. For Combination 3 clones (**Figure 2A and Supplementary Figure 5**), cells were inoculated to OD600 of 0.02 in YNB-glucose broth, grown overnight, washed using water, and inoculated into YNB-galactose broth to OD600 of 0.2 to induce ToxAmp. The cells were further sub-cultured in YNB-galactose broth. The yEGFP fluorescence was analysed using A BD FACSCanto II flow cytometer.

#### Testing the stability of yEGFP-expressing construct

For each subculture, the cells were inoculated to OD600 equal to 0.001 for YPD broth and 0.008 for YNB-glucose broth, grown overnight for yEGFP fluorescence measurement when cells were in the exponential growth phase, and grown to 24 hours to start a new culture. The yEGFP fluorescence was analysed using A BD FACSCanto II flow cytometer.

#### Testing the stability of AeBlue-expressing construct

Cells were used to inoculate YPD medium at a concentration of OD_600_IJ0.005 and grown in a 200 rpm and 30 °C incubator for 24 hours, and the cells were used to start a new culture in the same conditions. For each culture, the cells were diluted to OD_600_IJ∼ 0.001, and 50µL of this dilution was spread on YPD agar plates (three plates for each culture). Agar plates were grown in a 30 °C incubator for about 48 hours and AeBlue was allowed to mature at room temperature until one week after plates had been spread. Colonies were counted and photographed.

### Genome sequencing

Oxford Nanopore whole genome sequencing was performed using the methods described previously.^8, 21^ The whole genome DNA was extracted using MagAttract HMW DNA Kit (Qiangen) with a modified protocol. A Rapid Barcoding Kit (SQK-RBK114.24, Oxford Nanopore), a R10.4.1 flow cell (FLO-MIN114, Oxford Nanopore), and a MinION sequencing Device (Oxford Nanopore) were used in Nanopore DNA sequencing. A MinKNOW software (23.x, Oxford Nanopore) was used in data collection and base-calling. The .fastq files were merged using a MergeFasta software (DNA BASER, Heracle BioSoft SRL Romania) and deposited in Sequence Read Archive database under submission number SUB14057532. A Canu assembler (Galaxy Version 2.2+galaxy0), a Maker genome annotation pipeline (Galaxy Version 2.31.11+galaxy2), and a MiniMap2 (Galaxy Version 2.24+galaxy0) were used in genome assembly and analysis through a Galaxy server (Galaxy Australia).^60^ JBrowse2 (Desktop version v2.6.2) was used to display genetic features.^61^

## Author Information

### Corresponding Author

**Bingyin Peng**-ARC Centre of Excellence in Synthetic Biology, Australia; Centre of Agriculture and the Bioeconomy, School of Biology and Environmental Science, Faculty of Science, Queensland University of Technology, Brisbane, QLD, 4000, Australia; Australian Institute for Bioengineering and Nanotechnology (AIBN), The University of Queensland, Brisbane, QLD, 4072, Australia https://orcid.org/my-orcid?orcid=0000-0001-6918-6016;

### Authors

Zeyu Lu-ARC Centre of Excellence in Synthetic Biology, Australia; Centre of Agriculture and the Bioeconomy, School of Biology and Environmental Science, Faculty of Science, Queensland University of Technology, Brisbane, QLD, 4000, Australia; Australian Institute for Bioengineering and Nanotechnology (AIBN), The University of Queensland, Brisbane, QLD, 4072, Australia

Liam McDonnell-ARC Centre of Excellence in Synthetic Biology, Australia; Centre of Agriculture and the Bioeconomy, School of Biology and Environmental Science, Faculty of Science, Queensland University of Technology, Brisbane, QLD, 4000, Australia

Will Anderson-Australian Institute for Bioengineering and Nanotechnology (AIBN), The University of Queensland, Brisbane, QLD, 4072, Australia

Francisco Peralta-ARC Centre of Excellence in Synthetic Biology, Australia; Centre of Agriculture and the Bioeconomy, School of Biology and Environmental Science, Faculty of Science, Queensland University of Technology, Brisbane, QLD, 4000, Australia

Tyson Watkins-ARC Centre of Excellence in Synthetic Biology, Australia; Centre of Agriculture and the Bioeconomy, School of Biology and Environmental Science, Faculty of Science, Queensland University of Technology, Brisbane, QLD, 4000, Australia

Hafna Ahmed-CSIRO Environment, Black Mountain Science and Innovation Park, Canberra, ACT 2601, Australia

Carlos Horacio Luna-Flores-Centre of Agriculture and the Bioeconomy, School of Biology and Environmental Science, Faculty of Science, Queensland University of Technology, Brisbane, QLD, 4000, Australia

Thomas Loan-CSIRO Agriculture & Food, Black Mountain Science and Innovation Park, Canberra, ACT 2601, Australia

Laura Navone-Centre of Agriculture and the Bioeconomy, School of Biology and Environmental Science, Faculty of Science, Queensland University of Technology, Brisbane, QLD, 4000, Australia; Eden Brew Pty Ltd, Glenorie, NSW, 2157

Matt Trau-Australian Institute for Bioengineering and Nanotechnology (AIBN), The University of Queensland, Brisbane, QLD, 4072, Australia; School of Chemistry and Molecular Biosciences (SCMB), the University of Queensland, Brisbane, QLD, 4072, Australia

Colin Scott-CSIRO Environment, Black Mountain Science and Innovation Park, Canberra, ACT 2601, Australia

Robert Speight-ARC Centre of Excellence in Synthetic Biology, Australia; Centre of Agriculture and the Bioeconomy, School of Biology and Environmental Science, Faculty of Science, Queensland University of Technology, Brisbane, QLD, 4000, Australia; Advanced Engineering Biology Future Science Platform, Commonwealth Scientific and Industrial Research Organisation (CSIRO), Black Mountain, ACT, 2601, Australia

Claudia E. Vickers-ARC Centre of Excellence in Synthetic Biology, Australia; Centre of Agriculture and the Bioeconomy, School of Biology and Environmental Science, Faculty of Science, Queensland University of Technology, Brisbane, QLD, 4000, Australia

#### Conflicts of Interes

Laura Navone declares competing interests in Eden Brew (Australia). Others declare no competing interests.

## Supplementary Information

Supplementary Result 1. Brownish mutation caused by adenine deficiency in AeBlue-expressing cells. Supplementary code 1. calculating kernel density using R. Supplementary Figure 1. The first-generation constructs of toxin-antitoxin-driven gene amplification system and the yeast transformation results. Supplementary Figure 2. The correlation between yEGFP fluorescence and copy number for strain TU2E4A1 clones (Figure 1). Supplementary Figure 3. Genetic features at the *ho* locus in strain TU2E4A1 clone A7 (Figure 1). Supplementary Figure 4. Sanger sequencing results for dead-RelE-expressing plasmid pToxEXP4R. Supplementary Figure 5. Characterisation of the clones isolated from the transformation of toxin-antitoxin-driven gene amplification constructs (Figure 2A). Supplementary Figure 6. Genomic features at the *ho* locus in strain 7DU2E4CUP1 clone B4. Supplementary Figure 7. *HO* (YDL227C) reappearing in copper(II)-induced evolution culture of strain 7DU2E4CUP1A1 clone B4. Supplementary Figure 8. Tandem gene repeats of yEGFP-RelB modules in copper(II)-induced evolution culture of strain 7DU2E4CUP1A1 clone B4. Supplementary Figure 9. Copper(II)-induced evolution of strain 7DU2E4CUP1A1 clones (Figure 2: Combination 1). Supplementary Figure 10. The effects of galactose induction on the 7DU2E4GAL1A1 clones (Figure 2: Combination 3). Supplementary Figure 11. The stability of the multi-copy integration of the yEGFP expression cassettes via the RelE-RelB-driven gene amplification (complement to Figure 3). Supplementary Figure 12. The cultivation plates showing the brownish colony in the sub-cultures of strain 7DU2E4A12 clone G2. Supplementary Figure 13. Sub-cultivation and adenine deficiency of the brownish clone isolated from 7DU2E4A12 clone G2.

## Acknowledgement

This research was supported partially by the Australian Government through the Australian Research Council Centres of Excellence funding scheme (project CE200100029). The views expressed herein are those of the authors and are not necessarily those of the Australian Government or Australian Research Council. This work is supported by Galaxy Australia, a service provided by the Australian Biocommons and its partners.

