## Supplementary Information for "The integration of tandem gene repeats *via* a bacterial type-II toxin-antitoxin-mediated gene amplification (ToxAmp) system and stability visualisation in *Saccharomyces cerevisiae*"

### Supplementary Result 1. Brownish mutation caused by adenine deficiency in AeBlue-expressing cells

When subculturing the AeBlue-expressing strain Clone G2 (Figure 4), we observed a brownish colony on the plate (Supplementary Figure 12). We were interested in the causes for the brownish colour. We performed the subculturing experiment to examine its genetic stability, qPCR to evaluate the *HO* promoter copy number, and yeast colony PCR to analyse the genetic features at the *ho* locus (Supplementary Figure 13). Three types of colonies appeared during the subculturing, the colonies showing parental brown colour, the colonies showing the orange colour, and the colonies showing the bluish colour (Supplementary Figure 13A). The proportion of the orange colonies increased to ~ 10% after 45-generation subcultures. Blue colonies maintained at a low proportion. In an orange colony, the *HO* promoter copy-number decreased to one, and yeast colony PCR also showed the absence of RelB-AeBlue (Supplementary Figure 13B-C). We further validated that the brownish colony and its derivatives were adenine-deficient and adenine supplement could turn the colony colour back to blue (Supplementary Figure 13D). This colour changes are consistent with the phenotype of yeast adenine-deficient mutants (Srb 1958; Kokina et al. 2014).

### Supplementary code 1. calculating kernel density using *R*

YPD <- data.frame(read.csv(file=file.choose(), header = FALSE))

x <- 1

while(x < 25){

tempdata <- Filter(Negate(is.na),YPD[,x])

densityTemp <- density(tempdata, bw=0.01)

if(x ==1){

result <- cbind(densityTemp[[1]],densityTemp[[2]])

}

else{

resultTemp <- cbind(densityTemp[[1]],densityTemp[[2]])

result <- cbind.data.frame(result, resultTemp)

}

x <- x + 1

}

write.csv(result, "YPDwo result.csv")

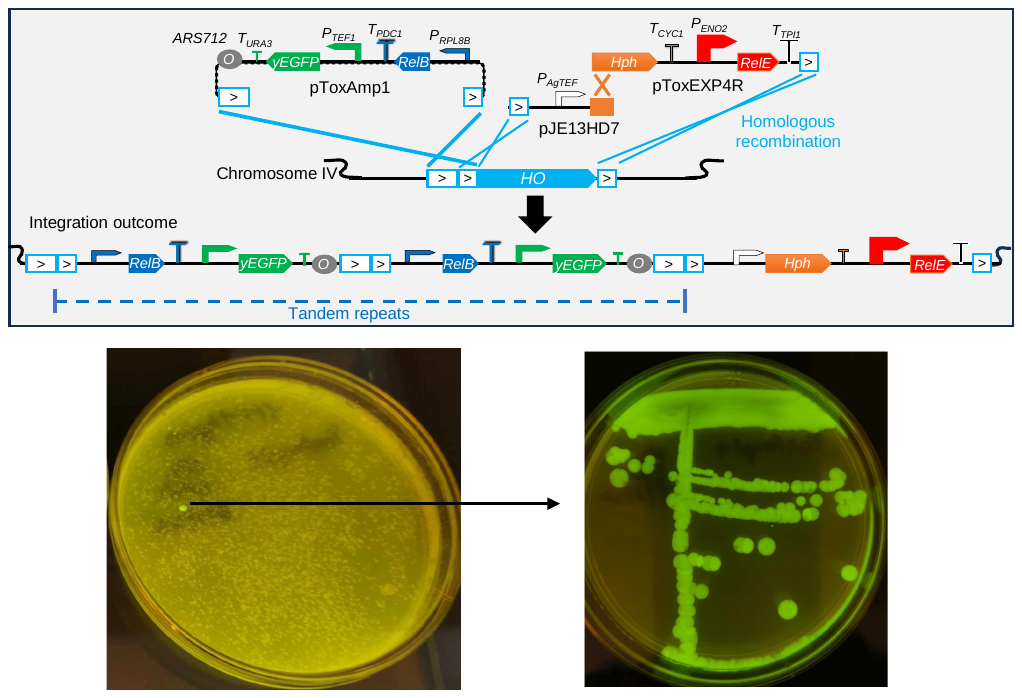

Supplementary Figure 1. The first-generation constructs of toxin-antitoxin-driven gene amplification system and the yeast transformation results. yEGFP fluorescence was imaged using a VWR Blue Light Transilluminator and a mobile phone camera.

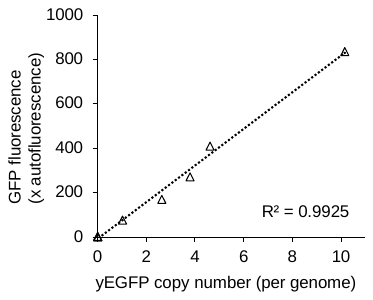

Supplementary Figure 2. The correlation between yEGFP fluorescence and copy number for strain TU2E4A1 clones (Figure 1).

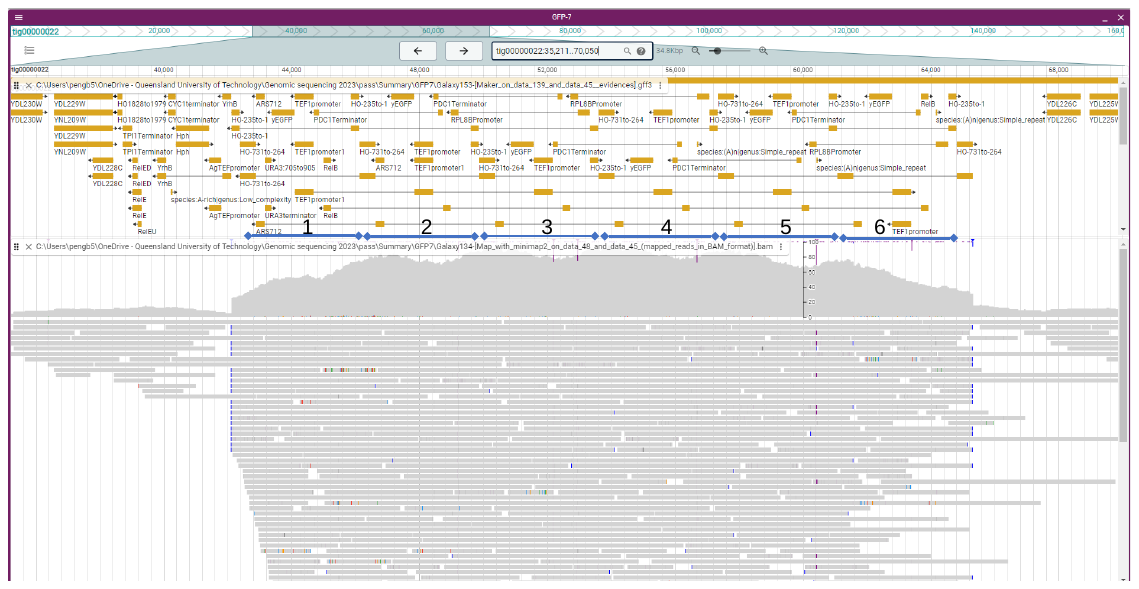

Supplementary Figure 3. Genetic features at the *ho* locus in strain TU2E4A1 clone A7 (Figure 1).

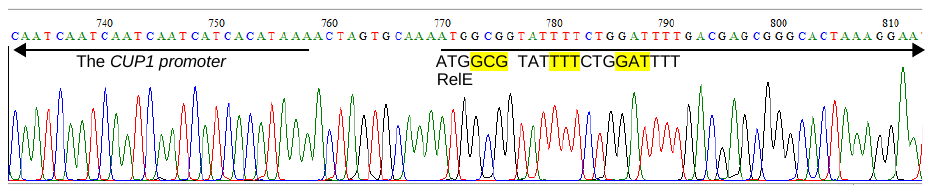

Supplementary Figure 4. Sanger sequencing results for dead-RelE-expressing plasmid pToxEXP4R.

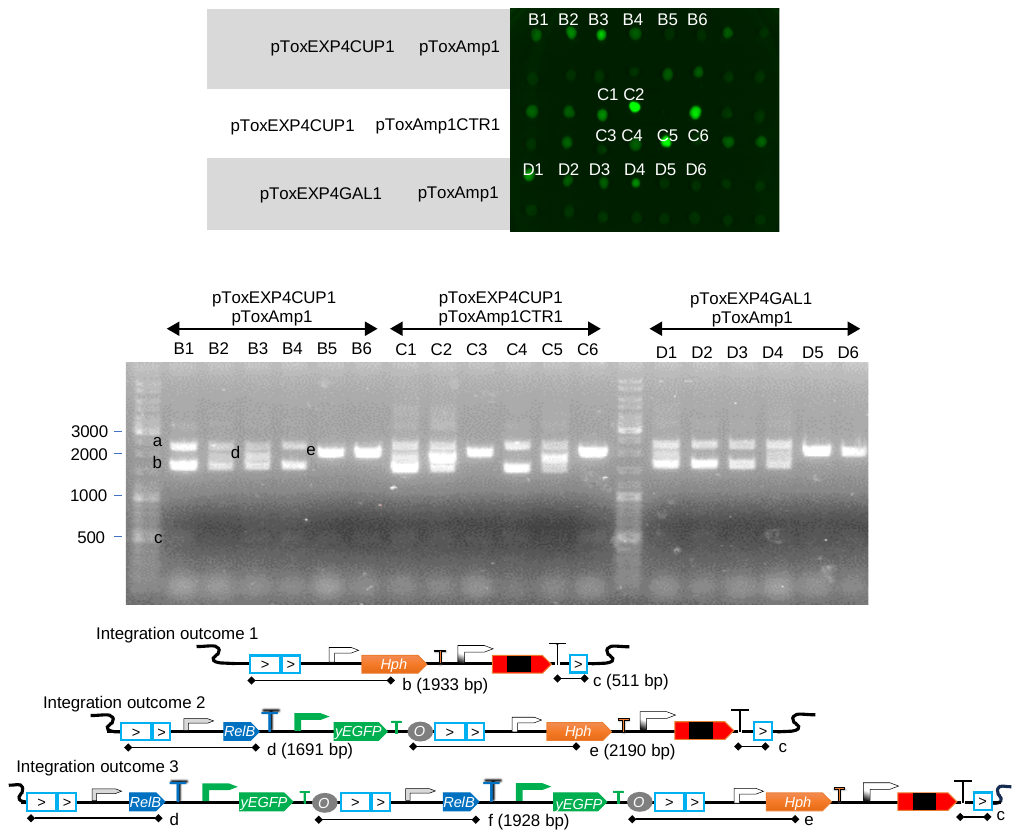

Supplementary Figure 5. Characterisation of the clones isolated from the transformation of toxin-antitoxin-driven gene amplification constructs (Figure 2A). The clones were replicated on YPD agar and imaged at Alexa 488 blot channel using a BioRad ChemiDoc^TM^ MP Imaging System. Yeast colony PCR was performed to confirm the integration outcomes (the diagram of the integration outcomes is same to that in Figure 2A).

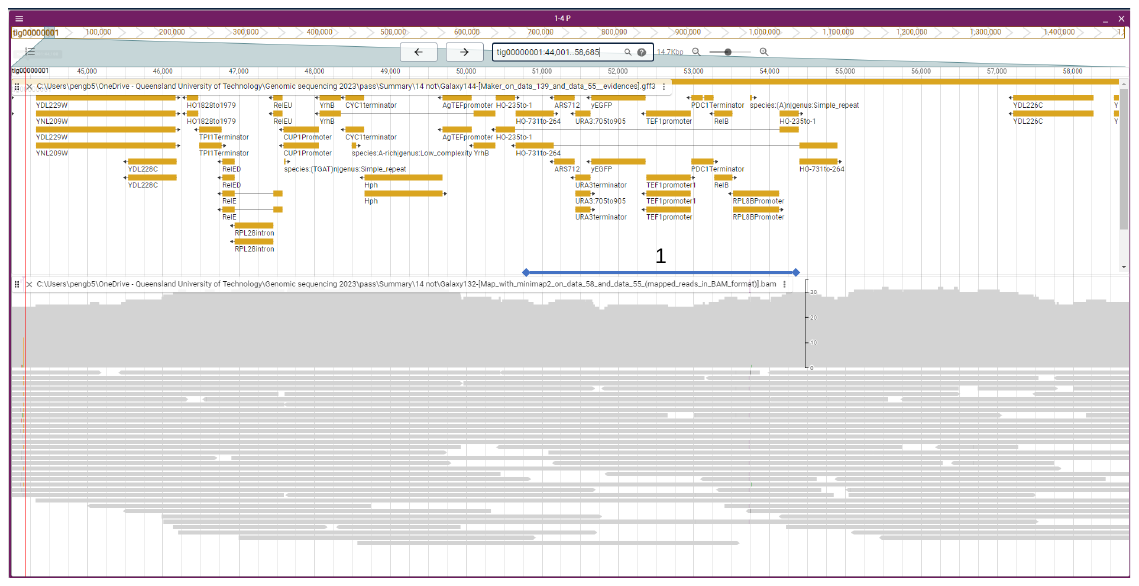

Supplementary Figure 6. Genomic features at the *ho* locus in strain 7DU2E4CUP1 clone B4.

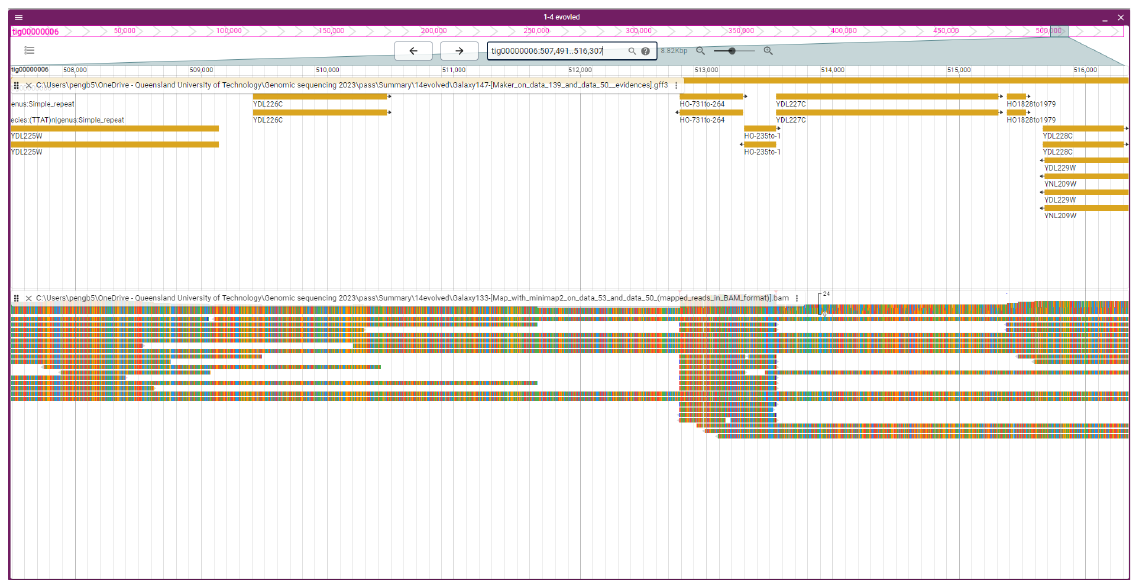

Supplementary Figure 7. *HO* (YDL227C) reappearing in copper(II)-induced evolution culture of strain 7DU2E4CUP1A1 clone B4.

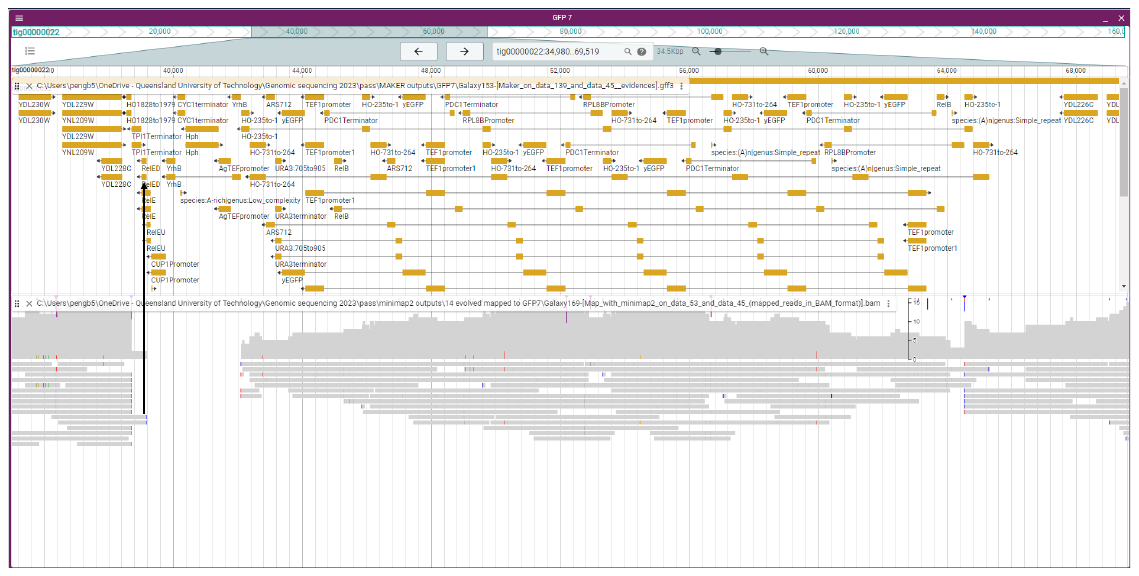

Supplementary Figure 8. Tandem gene repeats of yEGFP-RelB modules in copper(II)-induced evolution culture of strain 7DU2E4CUP1A1 clone B4. The reads were aligned to the assembly of strain TU2E4A1 clone A7 (Supplementary Figure 3).

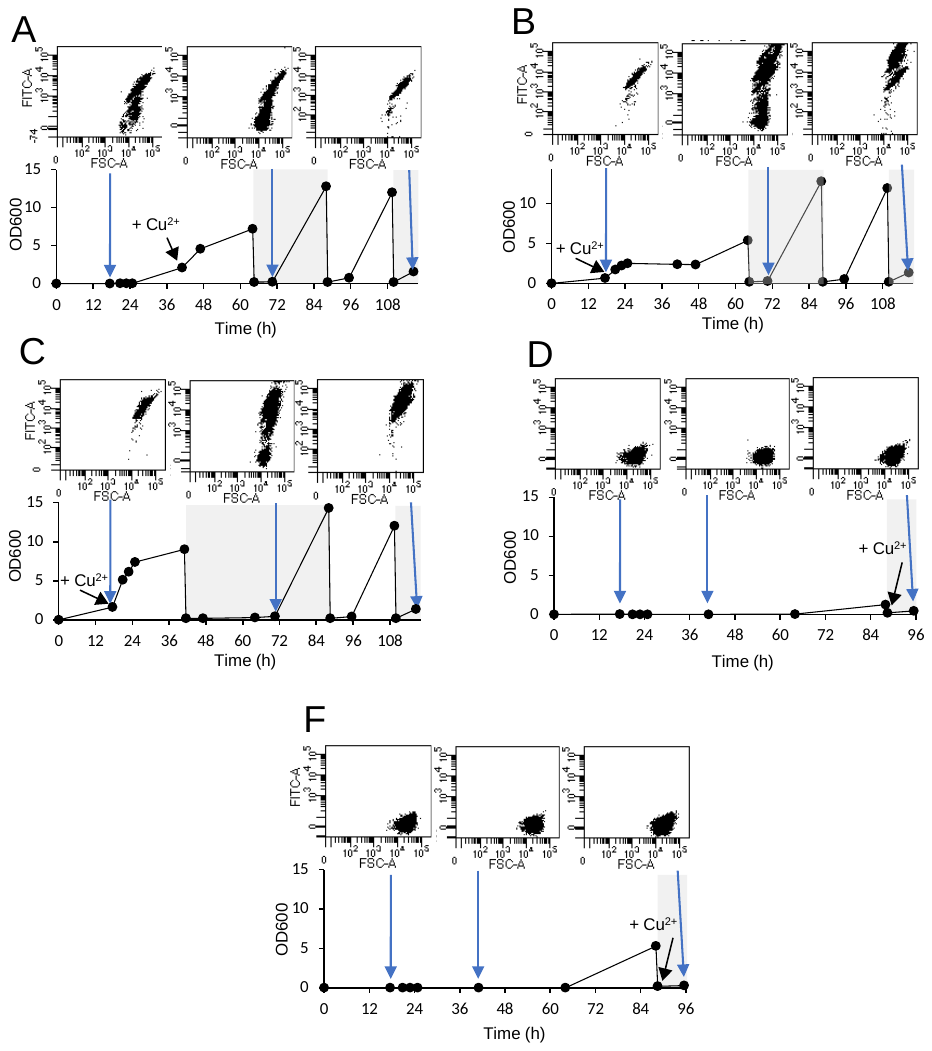

Supplementary Figure 9. Copper(II)-induced evolution of strain 7DU2E4CUP1A1 clones (Figure 2: Combination 1). Clones: B1 (A; single-copy integration), B2 (B; multi-copy integration), B3 (C; multi-copy integration), B5 (E; no integration of yEGFP-RelB module), and B6 (E; no integration of yEGFP-RelB module).

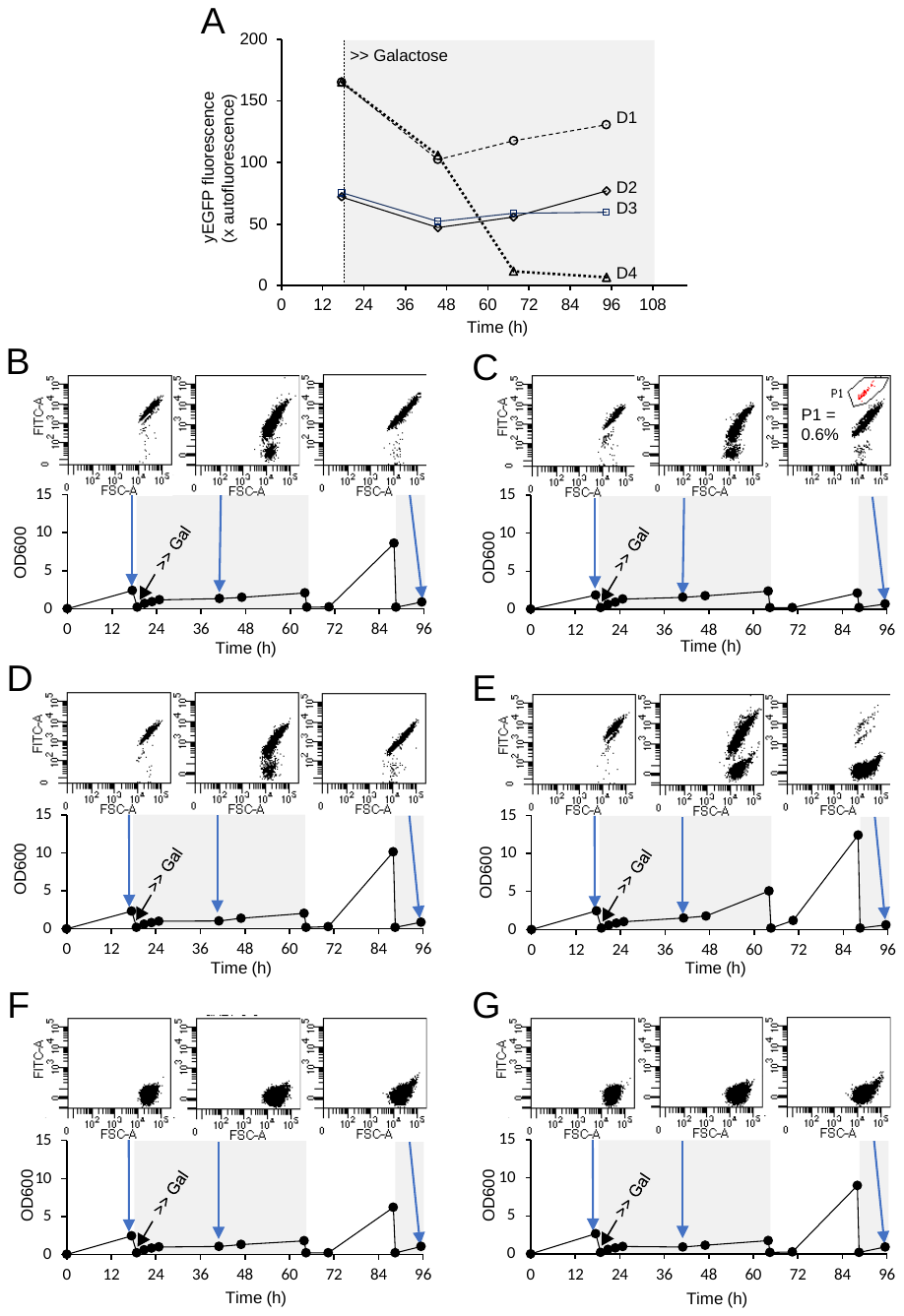

Supplementary Figure 10. The effects of galactose induction on the 7DU2E4GAL1A1 clones (Figure 2: Combination 3). (A) Overall yEGFP fluorescence levels. Clones: D1 (B; multi-copy integration), D2 (C; single-copy integration), D3 (D; single-copy integration), D4 (E; multi-copy integration), D5 (F, no integration of yEGFP-RelB module), and D6 (G, no integration of yEGFP-RelB module).

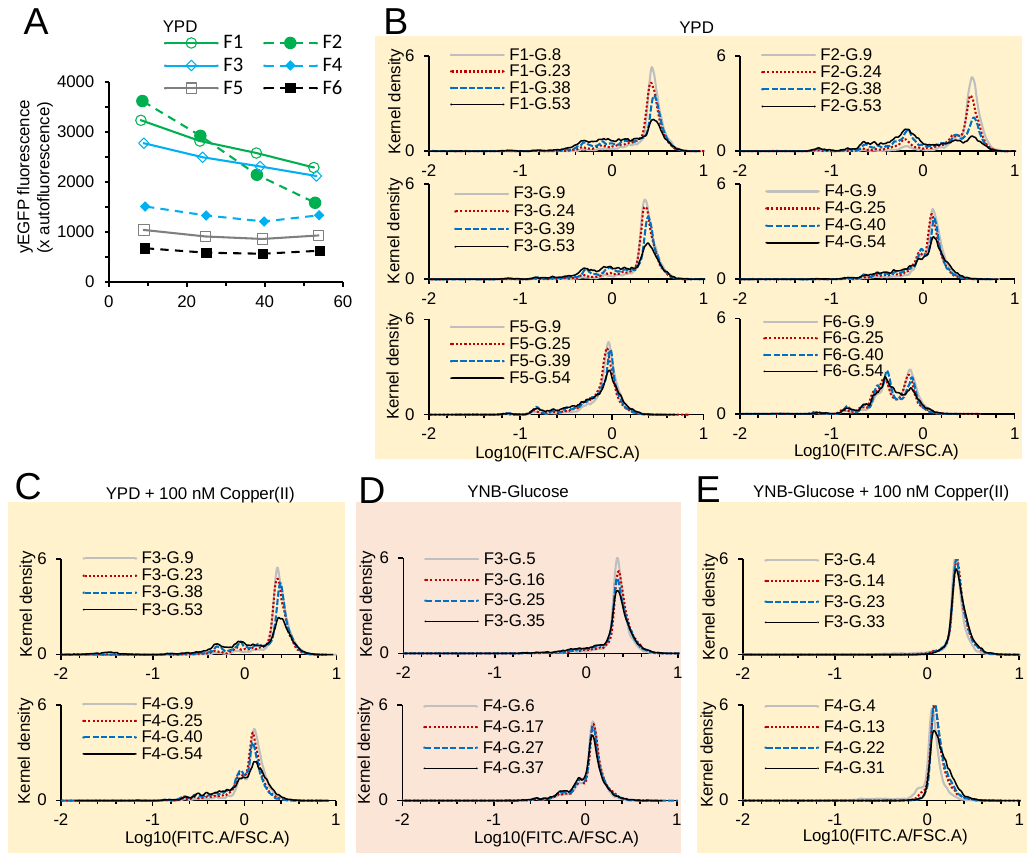

Supplementary Figure 11. The stability of the multi-copy integration of the yEGFP expression cassettes via the RelE-RelB-driven gene amplification (complement to Figure 3).

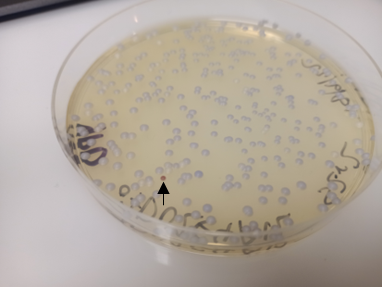

Supplementary Figure 12. The cultivation plates showing the brownish colony in the sub-cultures of strain 7DU2E4A12 clone G2.

Supplementary Figure 13. Sub-cultivation and adenine deficiency of the brownish clone isolated from 7DU2E4A12 clone G2. (A) Statistics of the colony colour in the subcultures of the original brownish clone. (B) The copy number of the *HO* promoter of the clone for each colour. Cells were sub-cultured in YPD broth and on YPD agar. Mean value ± standard deviation are shown; n = 3 technical replicates; 1 biological replicate for each colour was tested). (C) Yeast colony PCR showing the lost of RelB-yEGFP features in the orange colony (The PCR bands a, b, c, and d are shown in Figure 4A). (D) Plate dropping experiments showing the adenine deficiency in the brown isolate and its derivatives (n = 1). Yeast cells were re-suspended in water at OD600 ~ 1 and dropped on to YNB-glucose agar with serial dilutions. Adenine (100 mg L^-1^) was supplemented as indicated.
